# Authorship inflation and author gender in pulmonology research

**DOI:** 10.1101/446385

**Authors:** Blake Umberham, Caleb Smith, Nolan Henning, Matt Vassar

## Abstract

**Introduction:** Honorary authorship and equal gender representation are two pressing matters in scientific research. Honorary authorship is the inclusion of authors who do not meet the criteria established by the International Committee of Medical Journal Editors (ICMJE) authorship guidelines. The inclusion of honorary authors in the medical literature has led to an increase of the number of authors on studies and a decrease in single author studies in various fields.

**Methods:** Our primary objective was to assess authorship trends in two major pulmonology journals (selected on the basis of Google Scholar rankings): *Thorax* and *American Journal of Respiratory and Critical Care Medicine*. We reviewed all articles published in both journals in the years 1994, 2004, and 2014 using Web of Science and extracted data such as number of authors and gender of the first and last authors.

**Results:** The total number of authors steadily increased from 1994 to 2014. The median number of authors grew from about four in 1994 to nearly seven in 2014, which is approximately a 75% increase. When we compiled all the data, we found the percentage of female authors from both journals had increased from 17% to 29.9% during the study period.

**Discussion:** We found an increase in the average number of authors on pulmonology publications between 1994 and 2014 as well as an increase in the number of females with a lead or main author position. This may be due to a variety of factors, such as increased team science. However, our data in conjunction with data from other areas of medicine, indicate that honorary authorship may be contributing to the trends we identified.

## Introduction

Honorary authorship and equal gender representation are two pressing matters in scientific research. Honorary authorship is the inclusion of authors who do not meet the criteria established by the International Committee of Medical Journal Editors (ICMJE) authorship guidelines [1]. According to the ICMJE, each author should meet the following criteria: “1. Substantial contributions to the conception or design of the work; or the acquisition, analysis, or interpretation of data for the work; AND 2. Drafting the work or revising it critically for important intellectual content; AND 3. Final approval of the version to be published; AND 4. Agreement to be accountable for all aspects of the work in ensuring that questions related to the accuracy or integrity of any part of the work are appropriately investigated and resolved” [2 (page 2)]. The inclusion of honorary authors in the medical literature has led to an increase of the number of authors on studies and a decrease in single author studies in various fields [3–5]. Increasing authorship trends have been noted across many fields. For example, authorship inflation has been noted in journals pertaining to pharmacy [6], urology [7], radiology [8], orthopedic surgery [9], and neurology [10]. In high impact radiology publications, the number of authors per article doubled between 1980 and 2013 [8]. Furthermore, the average number of authors per article in orthopedic surgery journals has approximately doubled between 1958 and 2008 [9]. Despite these significant changes, authorship inflation is not the only issue clinical medicine literature is facing.

A second pressing matter in authorship is the disparity between male and female authors in medical research. Evidence suggests a gradual approach to equilibrium, with increased representation of female authors observed in surgical research [11], otolaryngology [12], dermatology [13], emergency medicine [14], and gastroenterology [15]. The number of female first authors on studies in three dermatology publications quadrupled over the last three decades [13], and the number of female first authors also quadrupled in otolaryngology studies between 1978 and 1998 [12]. However, these trends have not been confirmed in many fields of medical research.

Although many authorship studies have been performed throughout the medical literature, it has not been determined if these phenomena have impacted pulmonology research. We sought to determine whether the average number of authors and the representation of female authors have increased per report in pulmonology over the last twenty years. To accomplish this, we analyzed articles from two major pulmonology journals sampled at ten year intervals between 1994 and 2014.

## Methods

Prior to beginning our study, we conducted a review of the literature to determine whether or not a study had already been conducted over authorship patterns in the field of pulmonology. We also read publications pertaining to authorship in various fields of science to determine an appropriate number of journals to analyze and an approximation for the total number of articles to include in our study.

To determine the average number of authors and the percentage of female authors on pulmonology publications during the last 20 years, we analyzed two leading pulmonology journals: the *American Journal of Respiratory and Critical Care Medicine* (AJRCCM), and *Thorax*, selected based on Google Scholar Metrics h-5 index. We reviewed all articles published in both journals in the years 1994, 2004, and 2014, (n = 3,766) using Web of Science. We developed an abstraction manual with a corresponding Google Form that allowed us to list the name of the journal, year of publication, the names of the leading and main authors and their genders, the number of authors, the academic institution of the main author, and the Web of Science number for each article. Before data extraction, a training session was held to practice the process, address questions, and clarify the manual’s instructions.

In certain cases, the number of authors could not be determined due to corporate or anonymous authorship (n = 36). These cases consisted of eight editorials, seven guidelines, seven anonymous papers, four patient education handouts, four congresses, two reviews, one correction, one biography, one comment, and one research support article. Our final analysis of mean authorships per report equaled 3,730 articles. In the portion of our study pertaining the leading and main authors, we used each author as a data point (n = 7,532). We determined the genders of the leading and main authors from their first names and Internet research. In cases where the first names were not listed on the articles we analyzed, we searched Google Scholar and their academic institutions to determine the authors’ first names. In cases where we could not determine the gender of the leading and main authors based upon their first names and Internet searches, we used the *Baby Name Guesser* (formerly *Geoff’s Gender Guesser*) [16], to determine whether the name was more likely to belong to a male or a female. The Baby Name Guesser assigns a probability value of a name being male or female. If a name was at least twice as likely to be associated with a particular gender, the name was assigned to that particular gender. Probability values below two were considered undetermined. Ultimately, we excluded authors from our final analysis when gender was unable to be determined from their first name, internet search, or when results from the gender guesser were inconclusive (n = 498). A total of 7,034 authors comprised our final sample.

## Results

A total of 3,730 articles were reviewed for authorship inflation. The number of authors on articles published in *Thorax* between 1994 and 2014 generally increased. The median number of authors grew from about four in 1994 to nearly seven in 2014, which is approximately a 75% increase (**Figure 1**). The number of authors on articles published in the AJRCCM consistently exhibited growth between 1994 and 2014. Although the average number of authors did not increase significantly in this time period, the data clearly illustrate authorship inflation in 2014 compared with 1994 because the range and interquartile range (IQR) significantly increased (**Figure 2**). After compiling data from both journals, we found that the number of authors on articles published in both journals between 1994 and 2014 also increased. The average increased from about four authors for each article to about six authors for each article, which is approximately a 50% increase (**Figure 3**). In each case, the range and the IQR for the number of authors on each article continuously increased from 1994 to 2014 (**Table 1**). Despite an overall increase in the average number of authors on each paper, the mean for each data set did not consistently increase in each case. In addition, outliers in the data set resulted in inconsistent growth for the IQR and the range of the data.

**Figure 1.**
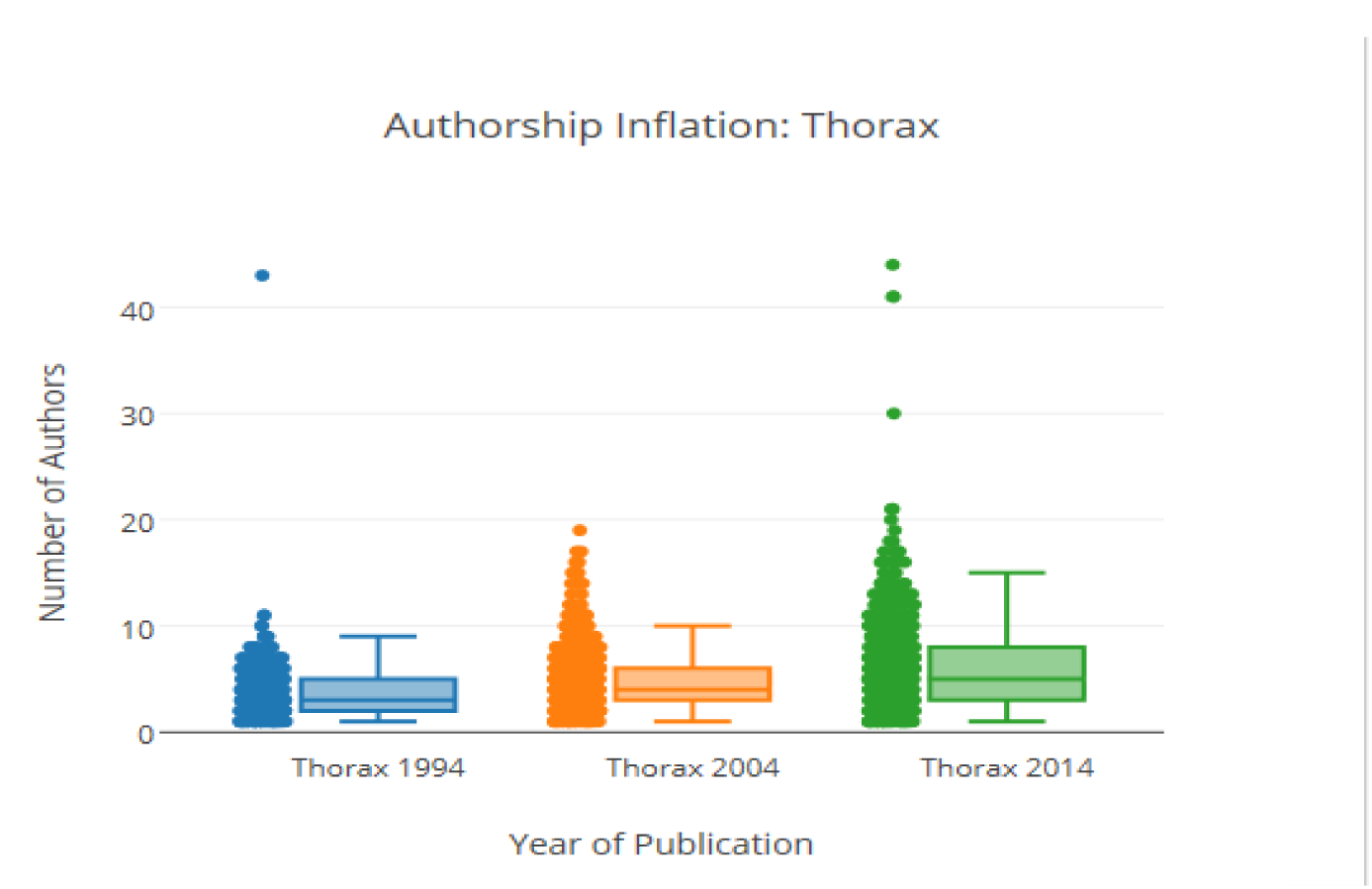

**Figure 2.**
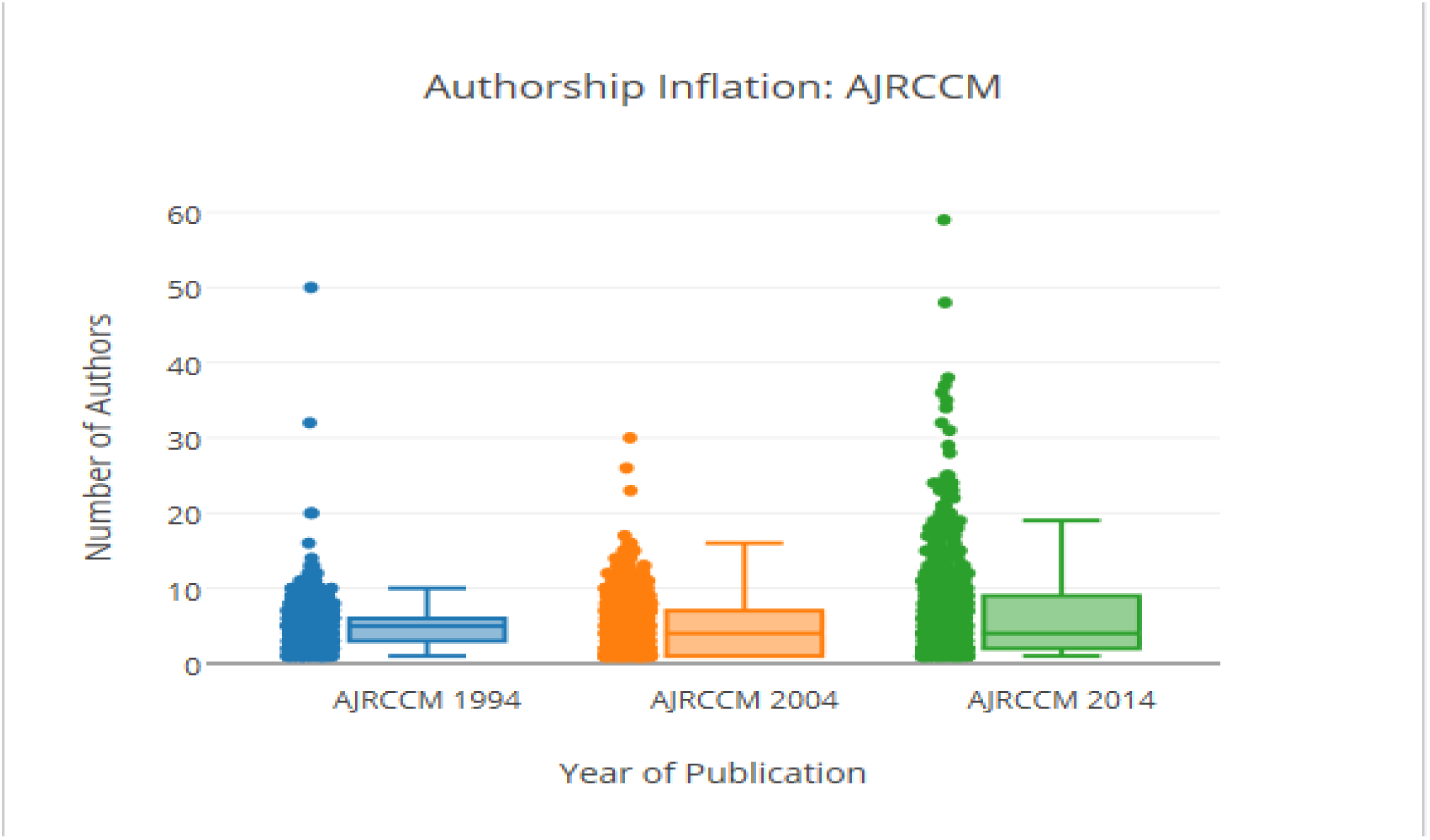

**Figure 3.**
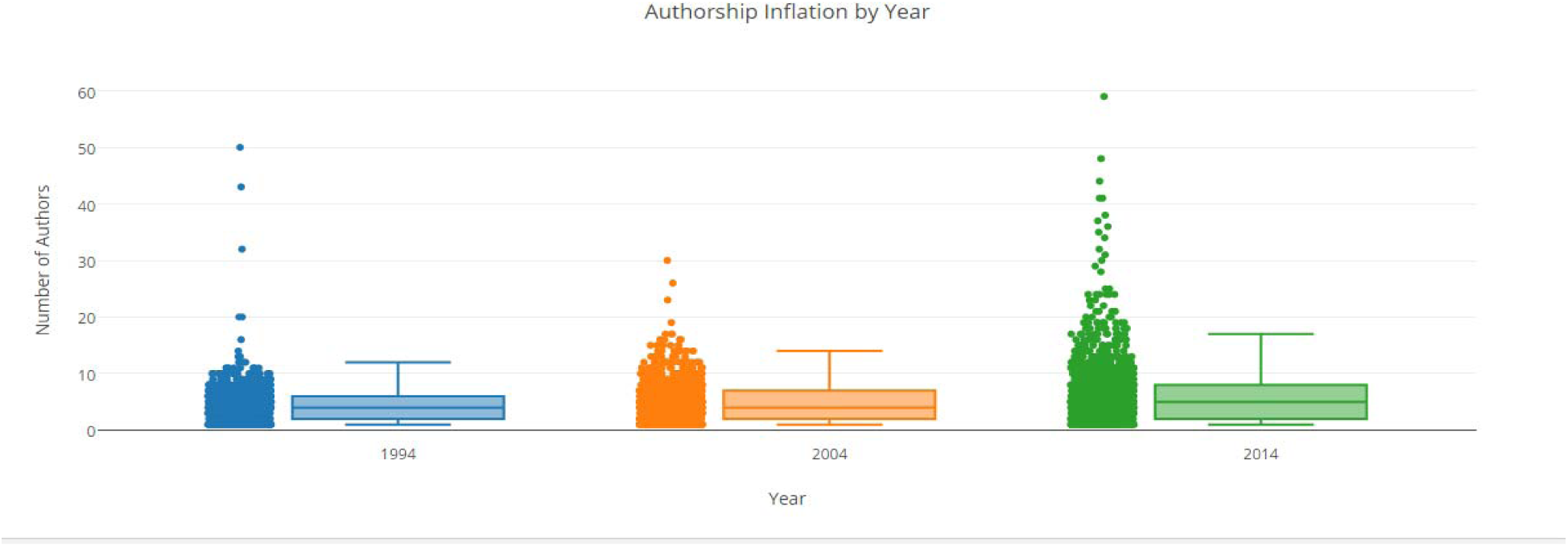

**Table 1.**

| Journal | Year | Mean | SD | Median | Q1 | Q3 | IQR | Range | Males | Females |
| --- | --- | --- | --- | --- | --- | --- | --- | --- | --- | --- |
| **Thorax** | 1994 | 4.074 | 7.28 | 3 | 1 | 5 | 4 | 129 | 82.90% | 17.10% |
|  | 2004 | 4.778 | 2.878 | 4 | 2 | 6 | 4 | 18 | 79.30% | 20.70% |
|  | 2014 | 6.016 | 4.394 | 5 | 3 | 7 | 4 | 43 | 70.50% | 29.50% |
| **AJRCCM** | 1994 | 5.256 | 8.532 | 5 | 3 | 6 | 3 | 205 | 83.10% | 16.90% |
|  | 2004 | 4.739 | 3.886 | 4 | 1 | 7 | 6 | 29 | 75.70% | 24.30% |
|  | 2014 | 6.442 | 6.591 | 4 | 2 | 8 | 6 | 58 | 69.70% | 30.30% |
| **Combined** | 1994 | 4.873 | 8.126 | 4 | 2 | 6 | 4 | 205 | 82.90% | 17.10% |
|  | 2004 | 4.759 | 3.406 | 4 | 2 | 6 | 4 | 29 | 79.30% | 20.70% |
|  | 2014 | 6.216 | 5.529 | 5 | 2 | 8 | 6 | 58 | 70.50% | 29.50% |

In respect to the gender question, we found that the percentage of female first and senior authors on pulmonology articles significantly increased between 1994 and 2014. In *Thorax*, the percentage of female authors increased from 16.9% in 1994 to 30.3% in 2014; in AJRCCM, the percentage of female authors increased from 17.1% to 29.5%, respectively. When we compiled all the data, we found the percentage of female authors from both journals had increased from 17% to 29.9% during the study period (**Table 1**).

## Discussion

We found an increase in the average number of authors on pulmonology publications between 1994 and 2014 as well as an increase in the number of females with a lead or main author position. The increase in the average number of authors on pulmonology publications parallels similar increases observed in pharmacy, urology, radiology, orthopedic surgery, and neurology journals [6–10]. In addition, the increase of female lead and main authors was also noted in surgical research, otolaryngology, dermatology, emergency medicine, and gastroenterology [11–15].

The increase in the average number of authors on publications in various clinical fields is notable because the trend could be unethical if individuals are offered authorship positions based on senior status or to increase a study’s likelihood for publication. If this is the case, then the number of authors per publication should be monitored and steps taken to minimize the negative effects and preserve the standing of authorship on a clinical publication. If this issue is not resolved, then the value of being an author on a publication is reduced. The increase in the number of female lead and main authors is notable because it provides evidence that females are becoming better represented in the medical community compared with previous decades.

The increase in the average number of authors on publications throughout various clinical fields could be caused by the inclusion of honorary and guest authors, a trend that has been noted in various high impact medical journals [17]. The inclusion of honorary and guest authors is most likely attributed to the importance placed on publications in the medical community. The publication of one’s work is necessary to progress through the medical field, which might prompt some researchers to seek unethical avenues to authorship. Furthermore, female authors have been underrepresented in the medical literature, which impedes their advancement through the medical community. The increase in females holding lead and main author positions could be a result of the government’s support of female researchers and students in the Science, Technology, Engineering, and Mathematics (STEM) initiatives [18] which led to the increasing number of females within the medical community [19].

Alternatively, the increase in the average number of authors on publications could be attributed to an increase in research complexity (such as multi-center clinical trials or epidemiological studies which requires specialized analytic knowledge); however, there is evidence to suggest that research complexity has not significantly impacted the increasing number of authors in medical publications [3]. The authorship inflation trend could also be influenced by the lack of awareness of ICMJE guidelines among researchers; however, this does not account for the authors’ personal beliefs about their contributions [20], which must be taken into consideration in determining the cause of authorship inflation since authors could feel that they must provide a collaborator with an authorship position despite the ICMJE guidelines. In addition to these causes, authorship positions could be assigned to satisfy requirements for clinical education hours and thereby contributing to unethical authorship claims in clinical publications. Despite the numerous unethical causes of authorship inflation, the phenomenon could be caused by the ease with which researchers can collaborate compared with twenty years ago. This allows for more individuals to contribute to a greater number of studies, signifying an ethical rise in authors per paper.

This study has several limitations. First, we analyzed only two pulmonology journals over twenty years; thus, our results might not be representative of all pulmonology publications. Second, we only analyzed articles from three years in the twenty year period; thus, our data does not reflect all aspects in the variation of the average number of authors on pulmonology publications. Third, research complexity, research quality, and geographical location were not accounted for in our study, which could have influenced the results of both portions of our study. Further research should be conducted in order to determine the specific causes of authorship inflation and whether or not authorship inflation is affecting all clinical fields at the same rate.

